# A novel allele in the *Arabidopsis thaliana* MACPF protein CAD1 results in deregulated immune signaling

**DOI:** 10.1101/2021.01.28.428637

**Authors:** Danalyn R. Holmes, Melissa Bredow, Kathrin Thor, Sydney A. Pascetta, Irina Sementchoukova, Kristen R. Siegel, Cyril Zipfel, Jacqueline Monaghan

## Abstract

Immune recognition in plants is governed by two major classes of receptors: pattern recognition receptors (PRRs) and nucleotide-binding leucine-rich repeat receptors (NLRs). Located at the cell surface, PRRs bind extracellular ligands originating from microbes (indicative of ‘non-self’) or damaged plant cells (indicative of ‘infected-self’), and trigger signaling cascades to protect against infection. Located intracellularly, NLRs sense pathogen-induced physiological changes and trigger localized cell death and systemic resistance. Immune responses are under tight regulation in order to maintain homeostasis and promote plant health. In a forward-genetic screen to identify regulators of PRR-mediated immune signaling, we identified a novel allele of the membrane-attack complex and perforin (MACPF)-motif containing protein CONSTITUTIVE ACTIVE DEFENSE 1 (CAD1) resulting from a missense mutation in a conserved N-terminal cysteine. We show that *cad1-5* mutants display deregulated immune signaling and symptoms of autoimmunity dependent on the lipase-like protein ENHANCED DISEASE SUSCEPTIBILITY1 (EDS1), suggesting that CAD1 integrity is monitored by the plant immune system. We further demonstrate that CAD1 localizes to both the cytosol and plasma membrane using confocal microscopy and subcellular fractionation. Our results offer new insights into immune homeostasis and provide tools to further decipher the intriguing role of MACPF proteins in plants.

## Introduction

Mechanisms to sense and respond to pathogens have evolved in all forms of life. Eukaryotic cells contain different types of receptors that detect the presence of microbes and activate immune signalling cascades resulting in cellular reprogramming. In plant cells, recognition of pathogens is achieved through two major groups of receptors: those on the cell surface and those located intracellularly (Jones and Dangl 2006; Couto and Zipfel 2016). Receptors at the cell surface are known as pattern recognition receptors (PRRs), which bind conserved features of entire groups of microbes known as microbe-associated molecular patterns (MAMPs). MAMPs are typically integral to microbial lifestyles and are therefore under strong selective pressure (McCann *et al.* 2012). Classic examples include bacterial flagellin and Elongation Factor-Thermo Unstable (EF-Tu) (Couto and Zipfel 2016). In the model plant *Arabidopsis thaliana,* flagellin (or the minimal epitope flg22) is recognized by its cognate PRR FLAGELLIN SENSING 2 (FLS2), while EF-Tu (or the minimal epitope elf18) is recognized by EF-Tu RECEPTOR (EFR) (Couto and Zipfel 2016). PRR surveillance can be thought of as detection of ‘non-self’ that can perceive both pathogenic and non-pathogenic microbes (Couto and Zipfel 2016). Endogenous ‘damaged self’ molecules, such as cell wall fragments or peptides released during cell damage are similarly recognized by PRRs (Segonzac and Monaghan 2019). For example, the peptide AtPep1 binds and activates integral membrane receptors PEP-RECEPTOR 1 (PEPR1) and PEPR2, triggering classical immune responses (Couto and Zipfel 2016). PRRs have been shown to function in complex with co-receptors or auxiliary proteins (Macho and Zipfel 2014; Couto and Zipfel 2016). One example of a co-receptor is BRASSINOSTEROID INSENSITIVE 1-ASSOCIATED KINASE1 (BAK1), that forms complexes with several PRRs including FLS2, EFR, and PEPR1/2. PRR complex activation triggers an influx of calcium ions, an apoplastic oxidative burst, and phosphorylation-mediated kinase cascades leading to transcriptional reprogramming and basal immune responses (Couto and Zipfel 2016).

In order to successfully colonize plants, adapted pathogens have evolved ways to secrete effector proteins into plant cells to suppress immune responses (Dou and Zhou 2012). In the evolutionary ‘arms race’ that exists between hosts and their pathogens, resistant plants have evolved intracellular receptors to intercept effectors and overcome disease. Most intracellular immune receptors contain an internal nucleotide-binding region (NB) and C-terminal leucine-rich repeats (LRR), preceded by an N-terminal domain that contains either a coiled-coil (CC) or Toll/Interleukin1-receptor (TIR) region (Jones and Dangl 2006). The family is known as NB-LRR receptors (or NLRs), which are typically classed as either CC-NB-LRRs (CNLs) or TIR-NB-LRRs (TNLs). NLRs of both classes scan for the presence of pathogenic effectors, either through direct protein binding or by ‘guarding’ the state of effector targets (van der Hoorn and Kamoun 2008). Activation of NLRs triggers the production of reactive oxygen species (ROS), activation of mitogen-activated protein kinases (MAPKs), and a sharp increase in the phytohormone salicylic acid (SA) that eventually results in a form of programmed cell death known as the hypersensitive response (HR) (Jones and Dangl 2006). Recent work has demonstrated that activated NLRs can form heteromeric ‘resistosomes’ similar to ‘inflammasomes’ in animals (Dangl and Jones 2019; Tian and Li 2020). For example, the CNL HOPZ-ACTIVATED RESISTANCE 1 (ZAR1) assembles into a pentameric ring-shaped complex predicted to facilitate lysis by forming pores in the plasma membrane (Wang *et al.* 2019a; b), and the TNLs RESISTANCE TO PERONOSPORA PARASITICA 1 (RPP1) and RECOGNITION OF XOPQ1 (ROQ1) assemble into tetrameric clover-shaped holo-NADases (Ma *et al.* 2020; Martin *et al.* 2020). This localized defense response is accompanied by the release of systemic signals that result in broad-spectrum resistance in distant tissues (Jones and Dangl, 2006).

While the HR can be effective at containing an infection, mechanisms that result in microbial death remain largely unknown. An increase in antimicrobial compounds such as ROS, SA, and nitric oxide hamper pathogen growth, but few molecular ‘agents of defense’ have been uncovered in plants. One component of the mammalian immune system employs pore-forming proteins to directly target and lyse microbial membranes to clear infection (Rosado *et al.* 2007). Examples include perforin (PF), which can form oligomeric rings in target membranes, as well as complement proteins C5, C6, C7, C8, and C9, which together form a heteromeric cylindrical ring known as the membrane attack complex (MAC) (Bayly-Jones *et al.* 2017). Interestingly, pore-forming proteins are also an important aspect of pathogen virulence (Rosado *et al.* 2007). The MACPF signature motif Y/S-G-T/S-H-X7-G-G, in which X can be any amino acid, is found in proteins across kingdoms, including plants (Ni and Gilbert 2017; Yu *et al.* 2020).

There are four MACPF-motif containing proteins encoded in the model plant *Arabidopsis thaliana.* Although two of these, CONSTITUTIVE ACTIVE DEFENSE 1 (CAD1) and NECROTIC SPOTTED LESIONS 1 (NSL1), have been studied genetically (Morita-Yamamuro *et al.* 2005; Noutoshi *et al.* 2006; Tsutsui *et al.* 2006; Asada *et al.* 2011; Fukunaga *et al.* 2017; Chen *et al.* 2020), the molecular function of Arabidopsis MACPF proteins remains elusive. In this study, we isolated a novel allele of *CAD1* from a forward-genetics screen (Monaghan et al., 2014) designed to identify loci important for immune homeostasis. We show that immune signalling is deregulated when CAD1 is unstable and provide evidence that the N-terminus of CAD1 is particularly important for its integrity. Using confocal microscopy and subcellular fractionation, we show that CAD1 localizes to both the cytosol and the plasma membrane. Overall, our work confirms and builds on earlier observations and provides new insight into the biological function of MACPF proteins in plants.

## Materials and Methods

### Plant Materials and Growth Conditions

*Arabidopsis thaliana* plants of the ecotypes Columbia-0 (Col-0) and Nossen-0 (No-0) were grown either on soil as one plant per pot in controlled environment chambers maintained at 20-22 °C with a 10-h photoperiod, or on 0.8 % agar Murashige and Skoog (MS) plates supplemented with vitamins and 1 % sucrose at 20-22 °C with a 16-h photoperiod. Isolation of *modifier of bak1-5* (*mob*) mutants and Illumina sequencing of bulked segregants was described previously (Monaghan *et al.* 2014; Etherington *et al.* 2014). The *bak1-5 mob4* mutants were purified by one backcross to *bak1-5.* Single *cad1-5* mutants were obtained by crossing *bak1-5 mob4* to Col-0. Null alleles *cad1-2* (GABI_192A09), *cad1-3* (GABI_385H08), and *nsl1-1* (PSH_21828) were obtained through the Nottingham Arabidopsis Stock Centre (NASC). Due to seedling necrosis, these lines had to be propagated as heterozygotes. Other mutant lines were described previously and genotyped to confirm homozygosity: *bak1-5* (Schwessinger et al. 2011), Col *eds1-2* (Feys et al. 2005), and *ndr1-1* (Century et al. 1997). Double mutants were obtained by crossing homozygous plants and genotyping F_2_ segregants by allele-specific PCR. Homozygous doubles were propagated and confirmed in the F_3_ generation. Transgenic lines were generated by floral dip using *Agrobacterium tumefaciens* strain GV3101 according to standard protocols (Bent 2006). Only transgenic plants with single inserts, as indicated by 3:1 segregation on selection plates in the T_2_, were used for further analysis. Homozygous lines were selected based on appropriate antibiotic or herbicide resistance in the T_3_ generation.

*Nicotiana benthamiana* plants were grown as one plant per pot in controlled environment chambers maintained at 20-22 °C with a 16-h photoperiod. Transient transformation using *Agrobacterium tumefaciens* strain GV3101 was conducted on plants that were between 5-and 6-weeks old as described previously (Monaghan *et al.* 2014). All constructs were co-infiltrated with the silencing suppressor P19. All germplasm, primers, and constructs used in this study are listed in Table S1.

### Molecular cloning

For trans-complementation of *bak1-5 mob4*, a Gateway-compatible full-length genomic *gCAD1* (*At1g29690*) fragment containing the native *pCAD1* promoter (1027 bp upstream of the translational start codon, containing three predicted W-box cis elements as described in (Tsutsui *et al.* 2006)) and including the endogenous stop codon was amplified from Arabidopsis Col-0 genomic DNA using Phusion *Taq* polymerase (New England Biolabs) and cloned into pENTR using the D/Topo kit (Invitrogen). This was followed by recombination into pGWB1 (Nakagawa *et al.* 2007) by LR Clonase II (Invitrogen). Additional clones harbouring *CAD1* or *NSL1* were designed and constructed for different end-uses as outlined in Table S1. All constructs were verified by Sanger sequencing (The Centre for Applied Genomics, Toronto; or The Genome Analysis Centre, Norwich).

### Immunity assays

Immunogenic peptides flg22, elf18, and AtPep1 were synthesized by EZ Biotech (Indiana USA). MAMP-induced ROS burst and seedling inhibition were performed as previously described (Bredow *et al.* 2019). Syringe-inoculations with *Pseudomonas syringae* pv. *tomato* (*Pto*) DC3000 were done as previously described (Monaghan *et al.* 2009).

### RNA extractions and real-time quantitative PCR

RNA extractions were either performed using TRI reagent (Sigma), as described previously (Monaghan *et al.* 2014), or using the Aurum Total RNA Mini Kit following manufacturer’s directions (Bio-Rad). cDNA was synthesized using Superscript III (Invitrogen) and quantitative PCR was performed on a CFX96 Touch Real-Time PCR Detection System (Bio-Rad) using Sso Advanced Universal SYBR Green Supermix (Bio-Rad) following MIQE guidelines (Bustin *et al.* 2009).

### Protein extractions and co-immunoprecipitation

Plant tissues were ground to a fine powder in liquid nitrogen and total protein was extracted as previously described (Monaghan *et al.* 2014). Protein concentration of the extracts was estimated based on tissue weight or normalized using Pierce Coomassie Protein Assay Kit (ThermoFisher Scientific). Co-immunoprecipitation (coIP) assays were performed using α-rabbit TrueBlot agarose beads (eBioscience) as previously described (Monaghan *et al.* 2014).

### Microsomal fractionation

Total microsome preparations were isolated as described previously (Takahashi *et al.* 2014). Briefly, approximately 40-50 g (fresh weight) of above ground biomass was collected from 5-week-old Arabidopsis plants grown on soil. Leaf tissue was cut into small pieces in 400-500 mL of pre-chilled homogenizing medium (0.5 M sorbitol, 50 mM MOPS-KOH (pH 7.6), 5 mM EGTA (pH 8.0), 5 mM EDTA (pH 8.0), 1.5 % (w/v) PVP-40, 0.5 % (w/v) BSA, 2.5 mM PMSF, 4 mM SHAM, 2.5 mM DTT). Tissue was then homogenized using a polytron generator (PT10SK, Kinematica Inc., Lucrene, Switzerland) and filtrates were collected by sieving the homogenate through Mericloth and removing cellular debris by centrifugation at 4 °C (10,000 × *g* for 15 min). Heavy membrane fractions were collected by ultracentrifugation at 231,000 × *g* for 50 min and resuspended in 1 mL of microsome suspension (MS) medium (10 mM KH_2_PO_4_/K_2_HPO_4_ (K-P) buffer (pH 7.8), 0.3 M sucrose) with a teflon-glass homogenizer. Samples were centrifuged again at 231,000 × *g* for 50 min and the pellet was suspended in MS-suspension medium (2 mL) prior to homogenization using an electric teflon-glass homogenizer. Soluble protein was collected by grinding ~500 mg of leaf tissue under liquid N_2_. Ground tissue was resuspended in 500 mL of native protein extraction buffer (10 mM Tris-HCl (pH 7.5), 25 mM NaCl) and incubated at 4 °C for 1 h, with shaking. Lysates were centrifuged at 12,000 × *g* for 10 min (4 °C) to remove cellular debris. Protein concentration was determined using Pierce Protein Assay Kit following the manufacturer’s instructions (ThermoFisher Scientific) and normalized to ~2 μg/uL.

### Immunoblots

Immunoblots were performed according to standard protocols using polyvinylidene difluoride membranes (Bio-Rad) blocked with 5% non-fat powdered milk in Tris-buffered saline containing 0.05-0.1% Tween-20. BAK1-FLS2 co-IP blots were probed with α-BAK1 (Schulze *et al.* 2010) or α-FLS2 (Gimenez-Ibanez *et al.* 2009) followed by α-rabbit-TrueBlot-HRP (eBioscience). CAD1 accumulation blots were probed with primary α-GFP (Roche) and secondary α-mouse-HRP (Sigma Aldrich). Subcellular fractionation blots were probed with primary α-H^+^-ATPase (Agrisera Antibodies) and secondary α-rabbit-HRP (Sigma Aldrich), or α-CFBPase (cytosolic fructose-1,6-biphosphatase) (Agrisera Antibodies) and secondary α-mouse-HRP (Sigma Aldrich). Blots were imaged by enhanced chemiluminescence (ECL) using Clarity Substrate (Bio-Rad) or ECL Prime (GE Healthcare) on a Chemidoc Imager (Bio-Rad) or using an X-Ray film developer.

### Confocal microscopy

Fluorescent proteins and dyes were excited using a 488 Argon laser and collected using separate tracks to detect different emission wavelengths; GFP or YFP emission was collected at 510-540 nm; SynaptoRed/FM4-64 at 635-680 nm; chlorophyll autofluorescence at 680-700 nm. Images were taken using the Zeiss LSM710 confocal microscope in the Biology Department at Queen’s University.

### Statistics

GraphPad Prism 8 was used to perform statistical tests on all quantitative data.

### Accession Numbers

CAD1 (At1g29690); NSL1 (At1g28380); BAK1 (At4g33430); EDS1 (At3g48090); NDR1 (At3g20600).

## Results and Discussion

### Isolation of a novel *CAD1* allele that affects its protein abundance

The C408Y variant of the co-receptor BAK1 resulting from the *bak1-5* mutation strongly impairs signalling mediated by multiple PRR receptor complexes (Roux *et al.* 2011; Schwessinger *et al.* 2011). Prolonged exposure to flg22, elf18, or AtPep1 results in impaired growth of wild-type seedlings, likely the result of sustained immune signalling over-ruling normal growth and development. However, seedling inhibition is partially blocked in *bak1-5* mutants (Roux et al. 2011; Schwessinger et al. 2011). Although strongly reduced, immune responses are not completely blocked in *bak1-5* mutants, providing a useful immuno-deficient background in which to perform a forward-genetic screen. To identify novel regulators of immunity, we previously isolated *modifier of bak1-5* (*mob*) mutants that restore immune signalling in *bak1-5* (Monaghan *et al.* 2014; Stegmann *et al.* 2017). Here, we describe *mob4*, which partially restores the flg22-, elf18-, and AtPep1-triggered ROS production and seedling inhibition in *bak1-5* (**Figures 1A, S1**).

**Figure 1:**
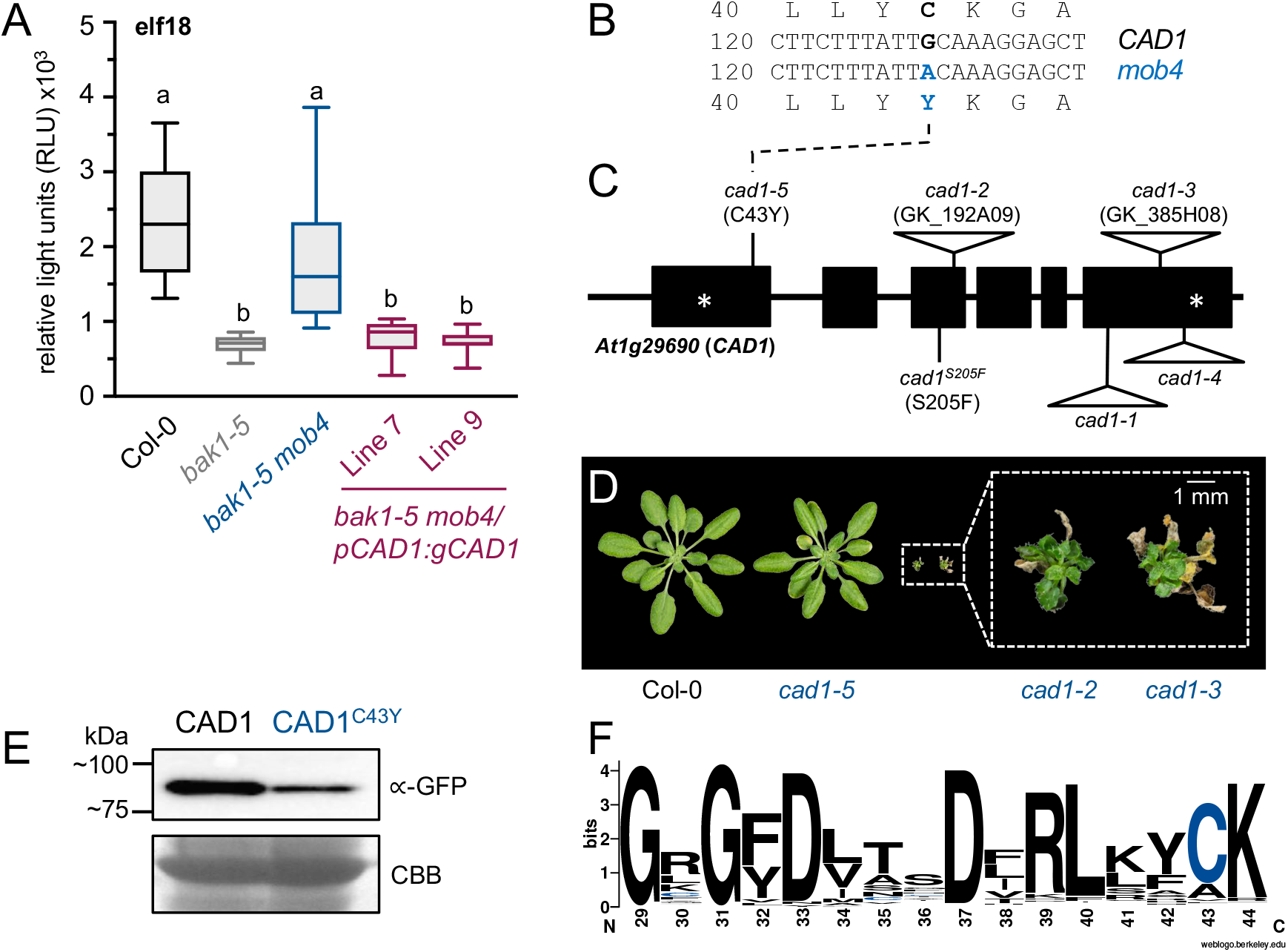
Isolation of a *CAD1* allele that affects its protein accumulation. **(A)** ROS production in the indicated genotypes after treatment with 100 nM elf18. Values are total photon count (relative light units) over 40 min, represented as box and whisker plots (n=6). Statistically significant (*p*<0.05) groups were analyzed by ANOVA followed by Tukey’s posthoc test and are indicated by lower-case letters. **(B)** Alignment comparing the amino-acid and codon sequences of CAD1 from Col-0 (top) and *mob4* (bottom), indicating the guanine-to-adenine mutation resulting in the C43Y protein variant. Numbers indicate the residue or open reading frame nucleotide position. **(C)** Schematic depicting the CAD1 gene, indicating the position of all published mutant alleles (the *cad1-1* and *cad1-4* alleles are described in (Morita-Yamamuro *et al.* 2005) and the *cad1*^*S205F*^ allele is described in (Chen *et al.* 2020)). Horizontal lines indicate untranslated regions and introns, and boxes indicate exons. The asterisks indicate the relative position of the start and stop codons. **(D)** Plants were grown in short-day conditions and photographed at 4 weeks post germination. Pictures were taken of plants grown together under identical conditions. **(E)** Western blot showing accumulation of CAD1-GFP following transient expression of *35S:CAD1-YFP* or *35S:CAD1*^*C43Y*^-*YFP* in *N. benthamiana.* Coomassie brilliant blue (CBB)-stained RuBisCO indicates relatively equal loading. **(F)** Consensus sequence of the newly-identified N-terminal motif found in MACPF proteins across the plant lineage, indicating strong conservation of CAD1^C43^, shown in blue. The Arabidopsis CAD1 sequence was used as a query in the Phytozome12 BLAST tool, resulting in the identification of 523 orthologs. Protein sequences were aligned using MUSCLE in MEGAX and the consensus sequence of the 16-amino acid motif was analyzed using Weblogo (Crooks *et al.* 2004).

F_1_ progeny from *bak1-5 mob4* backcrossed to *bak1-5* were insensitive to elf18 in seedling growth inhibition assays. Analysis of F_2_ progeny from the same cross indicated that *mob4* is caused by a single, recessive locus (177/799 plants had regained elf18-triggered seedling growth inhibition; *x*^2^=3.455; p=0.0631). To identify the causative mutation in *bak1-5 mob4*, we bulked the elf18-sensitive F_2_ progeny and sequenced the mutant genome using the Illumina HiSeq platform. We identified unique single nucleotide polymorphisms (SNPs) in the *bak1-5 mob4* genome by comparing the mutant genome to the parental *bak1-5* genome that we previously sequenced (Monaghan *et al.* 2014). We used the CandiSNP web application (Etherington *et al.* 2014) to plot the genomic positions of unique SNPs identified in *bak1-5 mob4* with an allele frequency over 75%. This analysis clearly indicated linkage on the top arm of Chromosome 1 (**Figure S2A**). We used a subset of these SNPs as markers and could further refine this region between 9.43 Mbp and 10.8 Mbp after conducting linkage analysis in individual F3 backcrossed lines. Contained within this region were single non-synonymous cytosine-to-thymine (or the complement; guanine-to-adenine) transitions in six genes (**Figure S2B**). In particular, we found a mutation causing a cysteine-to-tyrosine substitution at position 43 in the MACPF protein CAD1 (**Figure 1B**). To confirm that this mutation was causative of the *mob4* phenotype, we stably transformed *bak1-5 mob4* with full-length genomic *CAD1* and found that elf18-triggered ROS production (**Figure 1A**) was fully complemented in two independent homozygous *bak1-5 mob4/pCAD1:CAD1* lines. This confirmed that the mutant variant CAD1^C43Y^ is responsible for *mob4* phenotypes. We thus renamed this allele *cad1-5* in accordance with previously isolated *cad1* alleles (Morita-Yamamuro *et al.* 2005; Tsutsui *et al.* 2006; Asada *et al.* 2011; Chen *et al.* 2020).

Previous work demonstrated severe constitutive cell death in a complete loss-of-function *cad1-1* allele (Morita-Yamamuro *et al.* 2005). We obtained additional null alleles, *cad1-2* and *cad1-3,* and found that they similarly resulted in stunted growth and seedling lethality (**Figure 1C-D**). We noted that *bak1-5 cad1-5* plants have smaller rosettes than both Col-0 and *bak1-5* plants, and develop necrotic lesions, particularly in older leaves (**Figure S3A**). The degree of necrosis varied depending on environmental growth conditions, but *bak1-5 cad1-5* plants were always able to complete their life cycle and set plenty of seed. This led us to hypothesize that *cad1-* related growth defects and cell death might be suppressed by *bak1-5*, which could explain why we recovered a recessive *cad1* allele from the *mob* screen. To test this, we crossed *bak1-5 cad1-5* with Col-0 to isolate single *cad1-5* mutants. We found that *cad1-5* mutants looked phenotypically similar to *bak1-5 cad1-5* (**Figure S3A**). These results rather suggest that the CAD1^C43Y^ variant expressed in *cad1-5* (**Figure S3B**) is partially non-functional and does not cause as severe phenotypes as the complete loss of CAD1 observed in null mutants (**Figure 1D**).

Cysteines contain sulphur atoms in their R groups that are capable of forming disulphide bonds within a polypeptide chain and are therefore important for tertiary folding and protein stability. To test if the C43Y mutation affects CAD1 stability, we transiently expressed cauliflower mosaic virus (CaMV) *35S* promoter-driven *35S:gCAD1-GFP* and *35S:gCAD1*^*C43Y*^-*GFP* in *Nicotiana benthamiana* and found that while both variants migrated to the expected size of ~90 kDa in western blots, accumulation of CAD1^C43Y^ was clearly reduced (**Figure 1E**). This suggests that C43 is important for CAD1 stability, which likely explains the partial loss-of-function phenotypes in *cad1-5* compared to null alleles. Interestingly, a multiple sequence alignment of 523 MACPF-motif containing proteins encoded across land plants revealed that C43 is found within a conserved N-terminal motif G-x-G-F/Y-D-x-x-x-D-x-R-L-x-x-**C**-K (**Figure 1F**), underscoring the possibility that this residue may be important for the stability of MACPF-containing proteins more broadly.

### Reduced CAD1 accumulation correlates with enhanced immune signalling

Equipped with a genetic resource with which to assess the biological function of CAD1 without the complications associated with null alleles, we sought to assess immune responses in *cad1-5.* Being unable to perform assays alongside *cad1* null-mutants, we generated two independent *cad1-5/pCAD1:gCAD1* rescue lines to confirm that any phenotypes we observed were indeed caused by the *cad1-5* mutation (**Figure 2A**). Similar to what has been reported for *cad1* null alleles (Morita-Yamamuro *et al.* 2005; Asada *et al.* 2011), we found that *cad1-5* plants displayed heightened basal expression of the immune marker gene *PR1* (**Figure 2B**), which was complemented in the two *cad1-5/pCAD1:gCAD1* lines. We next assessed pattern-triggered immune responses and found that *cad1-5* displayed enhanced elf18-and AtPep1-triggered ROS production (**Figures 2C, S4A**) and seedling growth inhibition (**Figure 2D, S4B**), which were again fully complemented in the *cad1-5/pCAD1:gCAD1* lines. Notably, we also found that *cad1-5* seedlings display extreme cell death when grown in the presence of elf18 (**Figure S4C**), and develop necrotic lesions 24 h after infiltration with elf18 as adult plants (**Figure S4D**).

**Figure 2:**
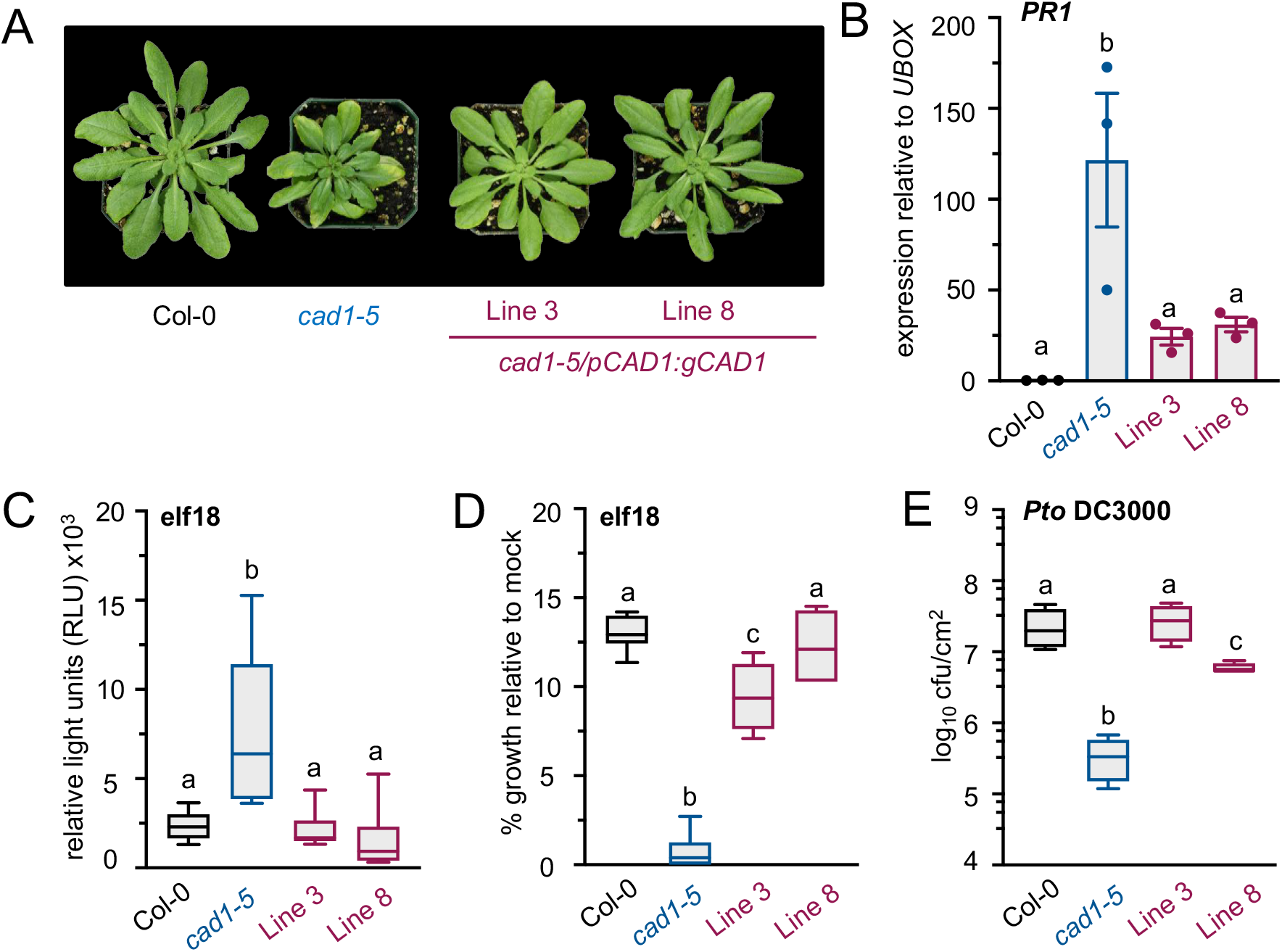
Enhanced MAMP-triggered responses and anti-bacterial immunity in *cad1-5*. **(A)** Plants were photographed after growth in a short-day chamber at 5 weeks post germination. **(B)** Expression of immune marker gene *PR1* is deregulated in *cad1-5* compared to Col-0 and complementing transgenic *cad1-5/pCAD1:gCAD1* lines. Real time quantitative reverse-transcription PCR of *PR1* was performed and plotted relative to expression of *UBOX*. Values are means +/− standard error from three independent experiments. **(C)** ROS production in the indicated genotypes after treatment with 100 nM elf18. Values are total photon count (relative light units) over 40 min, represented as box and whisker plots (n=6). **(D)** Seedling inhibition after 12 days of growth in 100 nM elf18 in the indicated genotypes, shown as % growth (fresh weight) relative to growth in control MS media (n=6). **(E)** Growth of the virulent bacterial pathogen *Pseudomonas syringae* pv. *tomato* strain DC3000 in the indicated genotypes 3 days after syringe-inoculation with 10^5^ colony forming units per millilitre (cfu/mL). Boxplots indicate log10 cfu/cm^2^ *in planta* bacterial growth (n=4, representing combined samples from 12 infected plants per genotype). All experiments were repeated at least three times with similar results. Statistically significant (*p*<0.05) groups were analyzed by ANOVA followed by Tukey’s posthoc test and are indicated by lower-case letters.

A previous study (Morita-Yamamuro *et al.* 2005) reported enhanced resistance to the virulent bacterial pathogen *Pseudomonas syringae* pv. *tomato* (*Pto* DC3000) in *cad1-1*; however, plants of this genotype display extreme necrosis, making interpretation of these data difficult. Follow-up experiments in which *CAD1* was transiently silenced using a dexamethasone-inducible construct confirmed that lower levels of *CAD1* result in enhanced resistance to *Pto* DC3000 (Asada *et al.* 2011). We also syringe-inoculated plants with *Pto* DC3000, and similarly found that *cad1-5* mutants harboured 10-fold less bacteria compared to Col-0 three days after inoculation, which was again complemented in the *cad1-5/pCAD1:gCAD1* transgenic lines (**Figure 2E**). To assess whether enhanced immune responses in *cad1-5* could be due to constitutive or enhanced PRR complex formation, we tested the flg22-induced association between the PRR FLS2 and its co-receptor BAK1. Co-immunoprecipitation (co-IP) assays using native α-BAK1 and α-FLS2 antibodies demonstrated similar complex formation in Col-0 and *cad1-5* (**Figure S4E**), indicating that autoimmunity in *cad1-5* is not caused by enhanced FLS2-BAK1 complex formation.

### Autoimmunity in *cad1-5* is *EDS1-*dependent

The null *cad1-1* mutant accumulates high levels of the immune hormone SA (Morita-Yamamuro *et al.* 2005). Transgenic expression of bacterial NahG, an enzyme that degrades SA, suppressed cell death in *cad1-1* (Morita-Yamamuro et al. 2005) – a classical hallmark of deregulated NLR-mediated signalling (Rodriguez *et al.* 2016). If CAD1 is an integral component of the plant immune response, it may be guarded by an NLR; inappropriate perturbation of CAD1 may thus activate NLR signalling resulting in uncontrolled cell death via the hypersensitive response (Rodriguez *et al.* 2016). NLRs can generally be classified into two groups: TNLs and CNLs (McHale *et al.* 2006), and it is possible to genetically manipulate components required for signalling through either class to delineate which type of NLR may be activated (Aarts *et al.* 1998). TNL signalling typically requires a family of nucleo-cytoplasmic lipase-like proteins including ENHANCED DISEASE SUSCEPTIBILITY 1 (EDS1), PHYTOALEXIN DEFICIENT 4 (PAD4) and SENESCENCE ASSOCIATED GENE 101 (SAG101) (Lapin *et al.* 2020), while CNL signalling typically requires the membrane-localized protein NON-RACE SPECIFIC DISEASE RESISTANCE 1 (NDR1) (Century *et al.* 1997; Coppinger *et al.* 2004). With this in mind, we generated *cad1-5 eds1-2* and *cad1-5 ndr1-1* double mutants to interfere with TNL- and CNL-triggered signalling pathways in *cad1-5*, respectively. We found that the small size and necrosis present in *cad1-5* plants was completely suppressed in *cad1-5 eds1-2* but not in *cad1-5 ndr1-1* (**Figure 3A**). Enhanced steady-state expression of *PR1* was similarly suppressed in *cad1-5 eds1-2,* but not *cad1-5 ndr1-1* (**Figure 3B**). We were next interested to test if the enhanced pattern-triggered responses and immunity in *cad1-5* were dependent on *EDS1*. Interestingly, although the heightened elf18-triggered ROS production (**Figure 3C**) was restored in *cad1-5 eds1-2,* elf18-triggered seedling growth inhibition and cell death was not restored in *cad1-5 eds1-2* (**Figure 3D**). In addition, we found that resistance to *Pto* DC3000 was suppressed in *cad1-5 eds1-2* (**Figure 3E**). Although enhanced susceptibility in *eds1-2* mutants (Aarts *et al.* 1998) could confound our interpretation, we propose that most of the enhanced immune phenotypes observed in *cad1-5* are dependent on *EDS1-*mediated signaling. EDS1 forms distinct signal-competent heteromeric complexes with SAG101 and PAD4 (Lapin *et al.* 2019). Interestingly, as previous work indicated that *pad4* was unable to suppress most *cad1-* related immune phenotypes (Tsutsui *et al.* 2008), likely also requiring helper NLRs such as N REQUIRED GENE 1 (NRG1) (Lapin *et al.* 2019, 2020). Altogether, these genetic analyses suggest that autoimmunity in *cad1-5* may be linked to activation of a TNL receptor; although this remains to be experimentally validated. Interestingly, *CAD1* was recently suggested to play a role in maintaining above-ground microbiota diversity, as *cad1* mutants present symptoms of dysbiosis (Chen *et al.* 2020). As high levels of SA in *cad1* may confound this interpretation, *cad1-5 eds1-2* might be an important tool to further dissect how CAD1 contributes to the maintenance of a healthy endophytic population.

**Figure 3:**
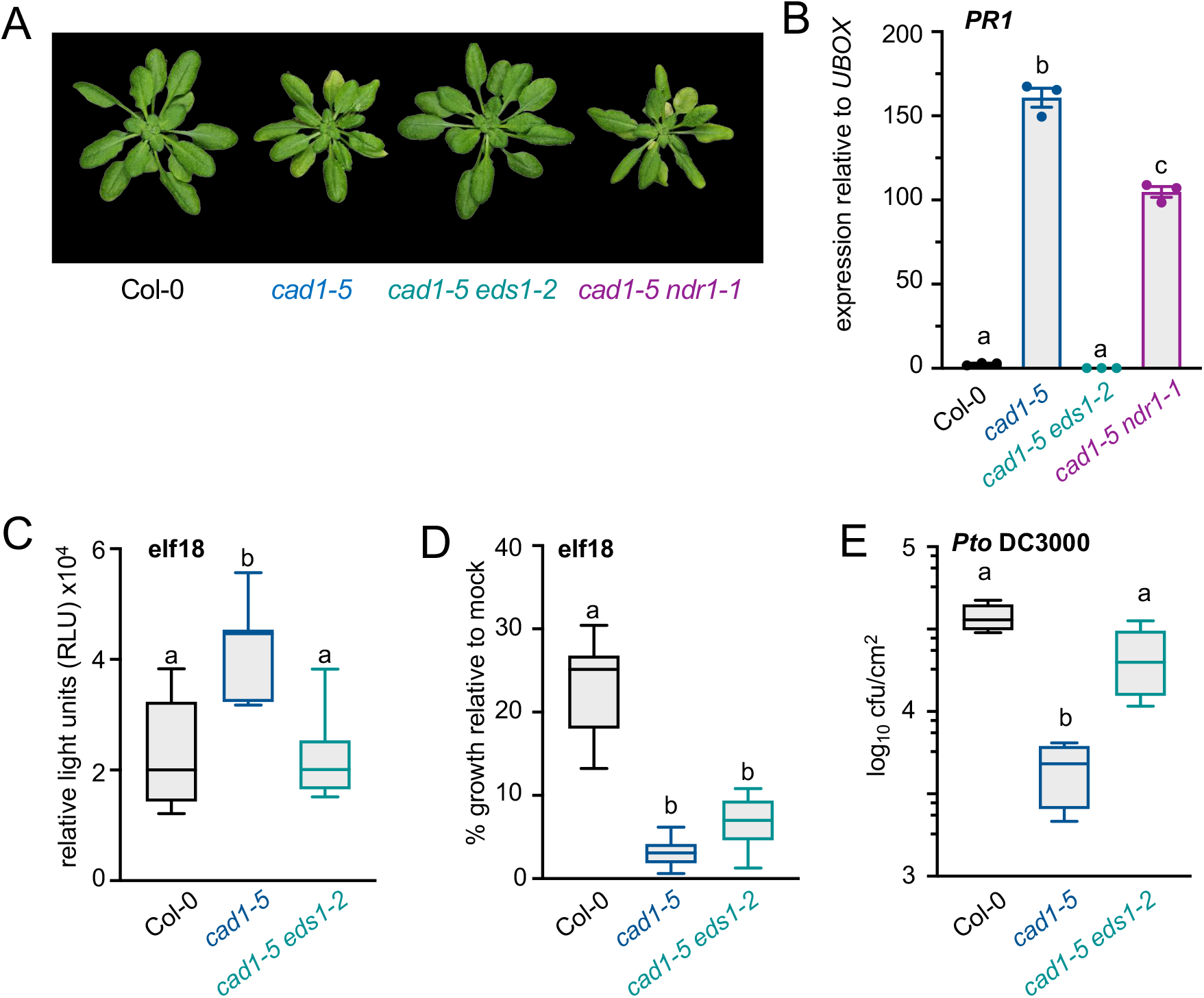
Autoimmunity in *cad1-5* is dependent on *EDS1.* **(A)** Smaller size and late-onset necrosis in *cad1-5* is suppressed by the introduction of *eds1-2*, which blocks signalling by TNL receptors, but not *ndr1-1*, which blocks signalling by CNL receptors. The indicated genotypes were generated by crossing, confirmed homozygous in the F_3_ generation, and photographed after growth in a short-day chamber at approximately 5 weeks post germination. **(B)** Real time quantitative reverse-transcription PCR of *PR1* was performed and plotted relative to expression of *UBOX*. Values are means + standard error from three independent experiments. **(C)** ROS production after treatment with 100 nM elf18. Values represent total photon count (relative light units) over 40 minutes (n=6). **(D)** Seedling inhibition after 12 days of growth in 100 nM elf18 in the indicated genotypes, shown as % growth (fresh weight) relative to growth in control MS media (n=6). **(E)** Growth of the virulent bacterial pathogen *Pseudomonas syringae* pv. *tomato* strain DC3000 in the indicated genotypes 3 days after syringe-inoculation with 10^5^ colony forming units per millilitre (cfu/mL). Boxplots indicate log10 cfu/cm^2^ *in planta* bacterial growth (n=4, representing combined samples from 12 infected plants per genotype). All experiments were repeated at least three times with similar results. Statistically significant (*p*<0.05) groups were analyzed by ANOVA followed by Tukey’s posthoc test and are indicated by lower-case letters.

CAD1 belongs to a small family of MACPF proteins in Arabidopsis. Included in this family are NSL1 (Noutoshi et al., 2006), and as-yet-uncharacterized proteins encoded by *At1g14780* and *At4g24290* (Morita-Yamamuro et al., 2005). Interestingly, loss-of-function *nsl1-1* mutants in the No-0 ecotype display spontaneous cell death similar to *cad1* mutants (Noutoshi *et al.* 2006). Null *nsl1-3* mutants in the Col-0 ecotype do not suffer from autoimmunity as in No-0, but do display runaway cell death when inoculated with the non adapted fungal pathogen *Colletotrihum orbiculare* or when treated with the MAMPs flg22 or elf18 (Fukunaga *et al.* 2017). Both autoimmune cell death in *nsl1-1* and induced cell death in *nsl1-3* are suppressed in *nsl1-1 eds16-1* (Noutoshi et al. 2006) and *nsl1-3 sid2* (Fukunaga et al. 2017), which cannot produce SA due to the loss of isochorismate synthase encoded by *SID2/ICS1/EDS16* (Wildermuth *et al.* 2001). Furthermore, hyperactive immunity is suppressed in *nsl1-3 pad4*, suggesting that NSL1 may also be guarded by a TNL (Fukunaga *et al.* 2017). Enhanced MAMP-induced immune responses in *nsl1* could also be suppressed by loss of *PENETRATION 2* (*PEN2*), a myrosinase involved in the synthesis of tryptophan-derived indole glucosinolates (Fukunaga *et al.* 2017). Future analysis of *cad1 pen2* mutants could determine if the loss of either or both CAD1 and NSL1 result in an increase of SA biosynthesis and tryptophan-derived secondary metabolites.

### CAD1 localizes to the plasma membrane and the cytosol

Perforin and complement proteins are targeted to non-autonomous cellular membranes to create pores and lyse those cells (Rosado *et al.* 2007). While it is possible that CAD1 and related proteins are deployed to pathogen membranes similar to their mammalian homologs, we could not detect secretory signal peptides in the amino-acid sequences of Arabidopsis MACPF proteins using the SignalP server (Emanuelsson *et al.* 2007). Previous work showed that NSL1 accumulates at the plasma membrane following MAMP induction in complementing, native-promoter driven and N-terminally tagged *nsl1-3/pNSL1:GFP-cNSL1* lines in the Col-0 ecotype (Fukunaga et al. 2017). During the course of our studies, we generated CaMV *35S* promoter-driven and C-terminally tagged *nsl1-1/35S:cNSL1-YFP* lines in the No-0 ecotype to similarly test sub-cellular localization of NSL1. Stunted growth and spontaneous necrosis in *nsl1-1* were complemented in two independent homozygous transgenic lines (**Figure S5A**), indicating functionality of the NSL1-YFP protein. We found that *35S*-driven NSL1-YFP localized to the plasma membrane even in the absence of MAMP-induction (**Figure S5B**), confirming membrane localization.

CAD1 has been identified in plasma membrane fractions in several independent proteomics studies (Alexandersson *et al.* 2004; De Michele *et al.* 2009; Elmore *et al.* 2012). To confirm the subcellular localization of CAD1, we transformed *cad1-5* mutants with *pCAD1:gCAD1-GFP* and tested genetic complementation of several *cad1-5* phenotypes in two independent homozygous lines. We found that expression of *pCAD1:gCAD1-GFP* complemented the *cad1-5* growth phenotype (**Figure 4A**) and enhanced expression of *PR1* (**Figure 4B**), suggesting that C-terminally tagged CAD1 is functional. Confocal microscopy suggested that CAD1-GFP localized to both the cytoplasm and the plasma membrane in *cad1-5/pCAD1:gCAD1-GFP* seedlings (**Figure 4C**). However, our micrographs were not always clear, probably due to the relatively low expression of native promoter-driven *CAD1*. Therefore, to confirm these results, we performed subcellular fractionation experiments using *cad1-5/pCAD1:gCAD1-GFP* plants to isolate soluble and microsomal fractions. Enrichment of plasma membrane in our microsomal preparations was confirmed by the clear presence of the membrane transporter H^+^-ATPase in western blots, while enrichment of cytosolic proteins in the soluble fraction was confirmed by probing for cytosolic fructose-1,6-biphosphatase (CFBPase). CAD1-GFP was clearly observed in both the microsomal and soluble fractions (**Figure 4D**), indicating that pools of CAD1 are found in both the cytosol and plasma membrane.

**Figure 4:**
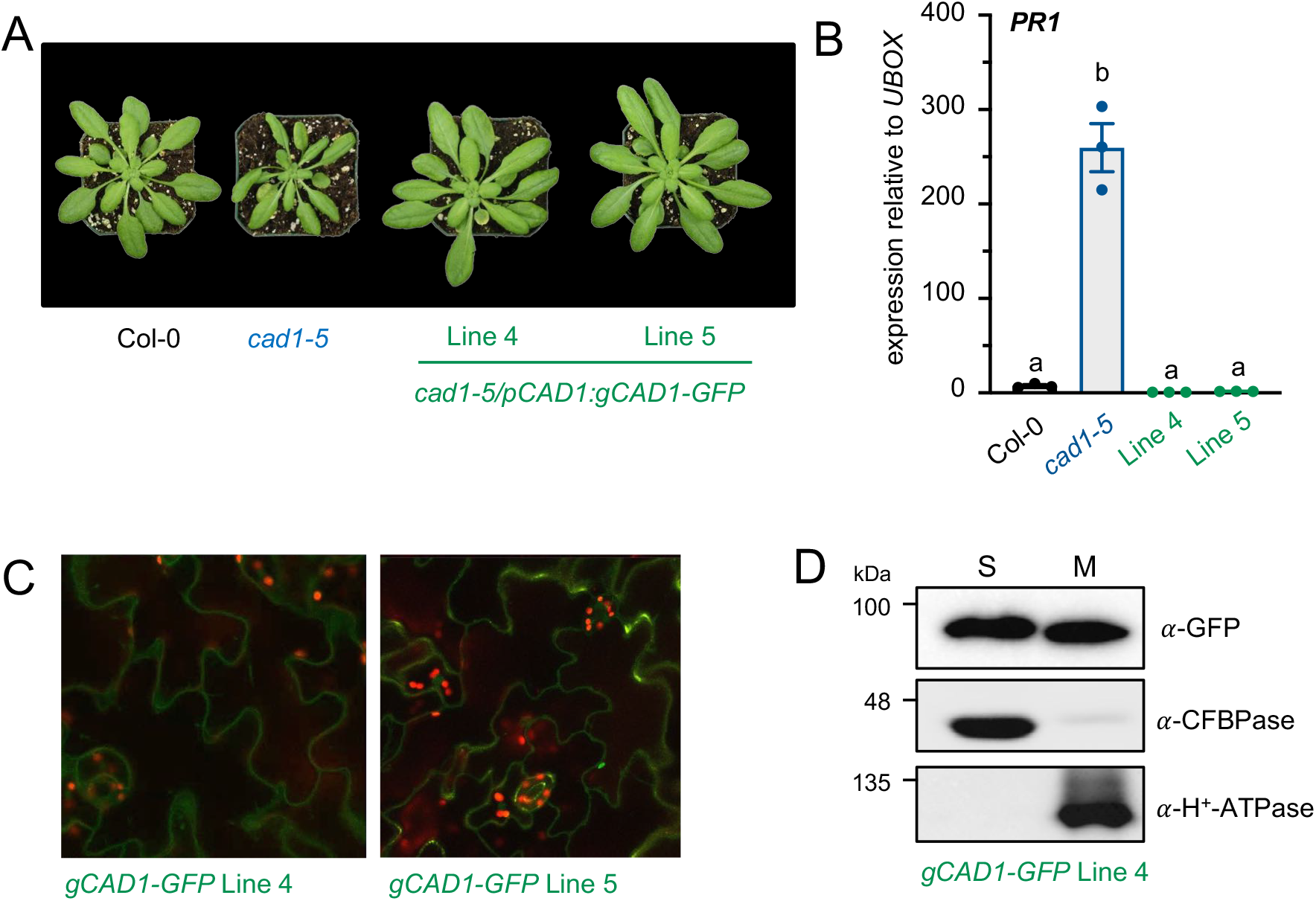
Subcellular localization of CAD1. **(A)** Plants were photographed after growth in a short-day chamber 5 weeks post germination. **(B)** Real time quantitative reverse-transcription PCR of *PR1* was performed and plotted relative to expression of *UBOX.* Values are means + standard error from three independent experiments. **(C)** Confocal microscopy images of *cad1-5/pCAD1:gCAD1-GFP* transgenic lines #4 and #5 with both the green and red channels overlaid; green indicates GFP fluorescence and red indicates auto-fluorescence from chloroplasts. **(D)** Western blot on soluble and microsomal protein fractions of *cad1-5/pCAD1:gCAD1-GFP* transgenic line #4. Detection of cytosolic CFBPase or membrane-bound H^+^-ATPase demonstrates successful fractionation. CAD1-GFP is clearly detected in both the soluble and microsomal fractions, indicating dual-localization.All experiments were repeated at least three times with similar results. Statistically significant (*p*<0.05) groups were analyzed by ANOVA followed by Tukey’s posthoc test and are indicated by lower-case letters.

During our studies, we also generated transgenic lines expressing N-terminally tagged *35S:GFP-cCAD1* in the *cad1-5* background. Of these, 18/22 T1 lines displayed severe necrosis and died on soil, resembling *cad1* null alleles. To test if N-terminally tagged CAD1 resulted in a dominant-negative variant, we transformed Col-0 with both *35S:GFP-cCAD1* and *35S:cCAD1.* Indeed, we found that 32/40 T1 lines of the genotype Col-0/*35S:GFP-cCAD1* displayed necrotic and small rosette size whereas the Col-0/*35S:cCAD1* control lines looked phenotypically similar to wild-type (**Figure S6**). These data suggest that the integrity of the N-terminus is important for CAD1 function, possibly by regulating its stability, localization, or binding partners. This seems to be in contrast to NSL1, which can be recombinantly tagged on either the N-(Fukunaga *et al.* 2017) or C-terminus (**Figure S5**) and still maintain function. Notably, a multiple sequence alignment of Arabidopsis MACPF proteins indicates variability in N-terminal residues (**Figure S7**), including an extended region of ~10 residues in CAD1.

Immunity is a complex trait involving many classes of proteins and layers of regulation. While proteins with MACPF motifs are known to function as pore-forming proteins in other organisms, their molecular function in plants remains unclear. Do these proteins form pores in membranes? And if so, how? Deploying pore-forming proteins as a defense mechanism would provide plants with a potent antimicrobial agent, and may therefore be targeted by pathogen effectors in the tug-of-war between host and microbe. This hypothesis could provide additional support to the suggestion that CAD1 and NSL1 are guarded by the immune system. Alternatively, these proteins may form pores in their own or neighbouring cell membranes, which could release DAMPs and potentiate immune signaling. Perforin contains both a membrane-associating ß-barrel region and membrane-inserting coiled-coil motifs, features that are essential for pore formation in mammalian MACPF domain proteins (Law et al., 2010). Neither CAD1 or NSL1 were predicted to contain any transmembrane regions using TMHMM Server v. 2.0; however, as both CAD1 and NSL1 localize to the cell membrane it is feasible that they form a heteromeric complex which together create a pore, similar to the complement proteins. In this scenario, it may be the integrity of the protein complex that is important for cell viability and the regulation of defense pathways. While this is supported by the finding that both *cad1* and *nsl1* mutants display similar (non-redundant) phenotypes, well-controlled protein-protein association studies are required to strengthen and test this hypothesis. We predict that uncovering the molecular function and biochemical mechanism of plant MACPF-containing proteins will be an exciting advance in plant cell biology.

## Supporting information

Holmes et al., Supplemental Data

## Acknowledgements

We thank two anonymous reviewers of an earlier version of this manuscript for their critical feedback that improved our study, and all members of the Monaghan and Zipfel laboratories for engaging discussions. We are grateful to Matthew Smoker and Jodie Pike (TSL Plant Transformation Facility) for generating several stable *Arabidopsis* transgenic lines. We thank the horticultural staff for growing plants and maintaining growth facilities at the John Innes Centre, as well as Jeffrey Rowbottom and Dale Kristensen for their assistance growing plants in the Queen’s Phytotron facility. We also thank Tony Papanicolaou (Queen’s University) for technical assistance with confocal microscopy. Peter Moffett (Sherbrooke University) provided the viral suppressor pBIN61:P19; and Darrell Desveaux (University of Toronto) provided *Pseudomonas syringae* pv. *tomato* DC3000.

## Author contributions

JM and CZ designed the study and funded the research; JM, DRH, MB, KT and SAP performed experiments; IS and KRS performed supporting experiments that are not shown. JM wrote the paper with input from all authors.

## Funding

This research was funded by grants from the Gatsby Charitable Foundation and the European Research Council (ERC) (‘PHOSPHinnATE’, Grant Agreement No. 309858) to CZ, as well as a Discovery Grant and Discovery Accelerator Supplement from the Natural Sciences and Engineering Research Council of Canada (NSERC), a John R. Evan’s Leader Fund Grant from the Canadian Foundation for Innovation (CFI) with matching funds from the Ontario Ministry of Research, Innovation and Science, and a Research Initiation Grant from Queen’s University to JM. JM was funded by a Long-Term Fellowship from the European Molecular Biology Organization (EMBO) during the early stages of this work. IS and KRS were funded by tandem NSERC Canada Graduate Scholarships for Master’s Students (CGS-M) and Ontario Graduate Scholarships (OGS), and MB was funded by an NSERC Postdoctoral Fellowship. No conflict of interest declared.

## Conflicts of interest

The authors have no conflicts to declare.

## Notes

### Competing Interest Statement

The authors have declared no competing interest.

