## Supplementary material for "A novel allele in the *Arabidopsis thaliana* MACPF protein CAD1 results in deregulated immune signaling": Holmes et al., Supplemental Data

##### Supplemental Figures

**Figure S1:** MAMP responsiveness is partially restored in *bak1-5 mob4*.

**Figure S2:** Identification of single nucleotide polymorphisms in *bak1-5 mob4* by whole-genome sequencing of bulked segregants.

**Figure S3:** *cad1-5* is not suppressed by *bak1-5*.

**Figure S4:** Enhanced MAMP-triggered responses in *cad1-5*.

**Figure S5:** NSL1 localizes to the plasma membrane.

**Figure S6:** N-terminally tagged CAD1 confers a dominant-negative effect.

**Figure S7:** Multiple sequence alignment of Arabidopsis MACPF proteins.

##### Supplemental Table

**Table S1:** Germplasm, primers, and constructs used in this study.

### Supplementary Figures

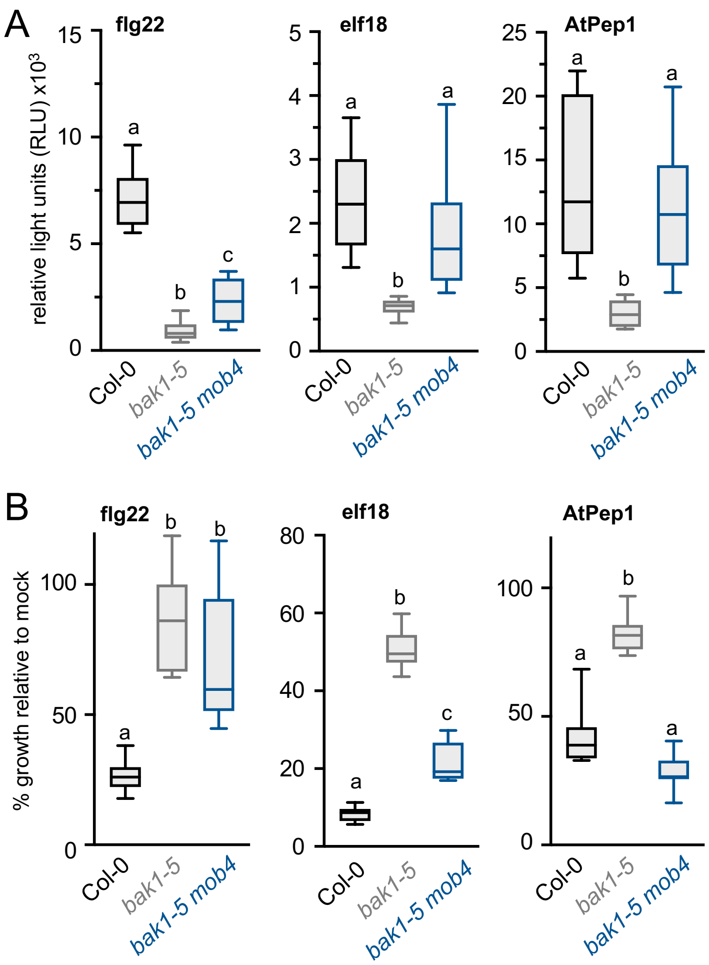


**Figure S1: MAMP responsiveness is partially restored in *bak1-5 mob4*.**
**(A)** ROS production in the indicated genotypes after treatment with 100 nM flg22, 100 nM elf18, or 1 µM AtPep1. Values represent total photon count (relative light units) over 40 min (n=8). **(B)** Seedling inhibition after 12 days of growth in 100 nM flg22, 100 nM elf18, or 1 µM AtPep1 in the indicated genotypes, shown as % growth (fresh weight) relative to growth in control MS media (n=12).
All experiments were repeated at least three times with similar results. Statistically significant ($p$<0.05) groups were analyzed by ANOVA followed by Tukey's posthoc test and are indicated by lower-case letters.

A

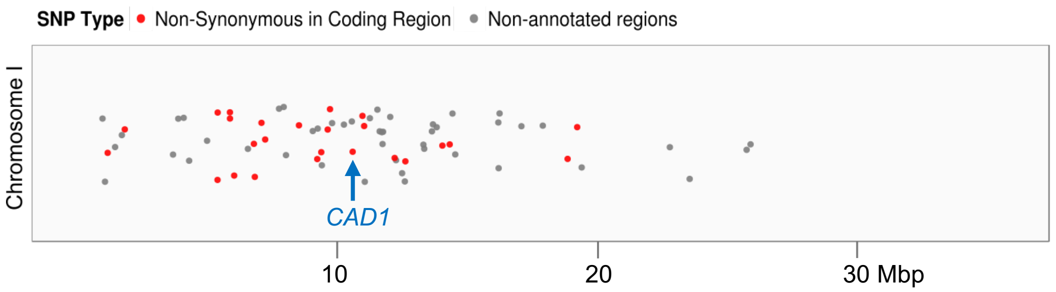


B

| Chr | Pos | Ref | Alt | Allele_Freq | Change | Gene ID | Gene Name | Sanger BC |
|---|---|---|---|---|---|---|---|---|
| 1 | 1116697 | C | T | 79.16 | S/F | AT1G04210 | Raf-related kinase | heterozygous |
| 1 | 1763961 | C | T | 86.2 | V/M | AT1G05840 | Proteolysis | heterozygous |
| 1 | 5271282 | C | T | 100 | S/N | AT1G15310 | SRP45 | heterozygous |
| 1 | 5276811 | C | T | 84.61 | R/K | AT1G15340 | MBD10 | heterozygous |
| 1 | 5740226 | C | T | 87.5 | R/K | AT1G16780 | Golgi proton pump | heterozygous |
| 1 | 5741941 | C | T | 85.18 | E/K | AT1G16780 | Golgi proton pump | heterozygous |
| 1 | 5901236 | C | T | 100 | S/F | AT1G17255 | Antisense gene | background |
| 1 | 6646943 | C | T | 86.2 | L/F | AT1G19230 | RbohE | heterozygous |
| 1 | 6681734 | C | T | 86.66 | E/K | AT1G19330 | Protein of unknown function | homozygous |
| 1 | 6934331 | C | T | 85 | T/I | AT1G19980 | Cytomatrix protein related | heterozygous |
| 1 | 7073500 | T | G | 75 | T/P | AT1G20400 | Protein of unknown function | not found |
| 1 | 8351039 | C | T | 92.3 | S/F | AT1G23550 | SRO1 | homozygous |
| 1 | 9042744 | C | T | 95.23 | E/K | AT1G26150 | PERK10 | homozygous |
| 1 | 9194612 | C | T | 93.93 | S/F | AT1G26610 | Nucleic acid binding | homozygous |
| 1 | 9437278 | C | T | 96.29 | T/I | AT1G27170 | TM receptor | homozygous |
| 1 | 9525661 | C | T | 100 | G/E | AT1G27430 | GYF protein | homozygous |
| 1 | 10381734 | C | T | 100 | C/Y | AT1G29690 | CAD1 | homozygous |
| 1 | 10752574 | C | T | 86.11 | P/S | AT1G30420 | ATP cassette | heterozygous |
| 1 | 10815851 | C | T | 90 | T/I | AT1G30530 | UGT78D1 | homozygous |
| 1 | 11969678 | C | T | 75 | E/K | AT1G33050 | Protein of unknown function | homozygous |
| 1 | 12375231 | C | T | 83.33 | L/F | AT1G34040 | Pyridoxal phosphate transferase | homozygous |
| 1 | 13774328 | C | T | 75 | A/V | AT1G36520 | Transposable element gene | not tested |
| 1 | 14051638 | C | T | 77.77 | S/F | AT1G37030 | Transposable element gene | not tested |
| 1 | 18510852 | C | T | 76.19 | T/I | AT1G49980 | DNA repair | homozygous |
| 1 | 18872523 | C | T | 76 | H/Y | AT1G50920 | GTP binding | homozygous |
| 3 | 13080899 | G | A | 76 | D/N | AT3G32091 | Transposable element gene | background |
| 3 | 13081217 | G | A | 87.5 | A/T | AT3G32091 | Transposable element gene | background |
| 4 | 7211101 | C | T | 76.47 | R/C | AT4G12030 | BASS5/BAT5 | background |
| 5 | 5599873 | G | A | 83.33 | G/S | AT5G17020 | XPO1 | not tested |
| 5 | 9186001 | G | A | 83.33 | G/R | AT5G26236 | Transposable element gene | background |
| 5 | 20881895 | G | A | 90.9 | R/* | AT5G51410 | LUC7 | heterozygous |
| 5 | 21236328 | G | A | 85 | P/L | AT5G52290 | SHOC1 | heterozygous |

**Figure S2: Identification of single nucleotide polymorphisms in *bak1-5 mob4* by whole-genome sequencing of bulked segregants.**
**(A)** CandiSNP output indicating the position of unique single nucleotide polymorphisms (SNPs) in bulked *bak1-5 mob4* mutants after a single cross to *bak1-5*. Red spots indicate SNPs that cause non-synonymous changes, while grey spots represent SNPs in regions that are not annotated as protein-coding. **(B)** Sequencing results of SNPs causing non-synonymous changes (red spots in A), indicating the Chromosome (Chr), Position (Pos), reference (Ref) and alternate (Alt) alleles, the % frequency the alternate allele was represented in Illumina reads (Allele_Freq), the expected amino acid change (Change), the gene ID, and the gene name. These SNPs were genotyped in individual recombinants isolated from a backcross (BC) by Sanger sequencing. The grey area indicates SNPs that were homozygous in each individual recombinant tested.

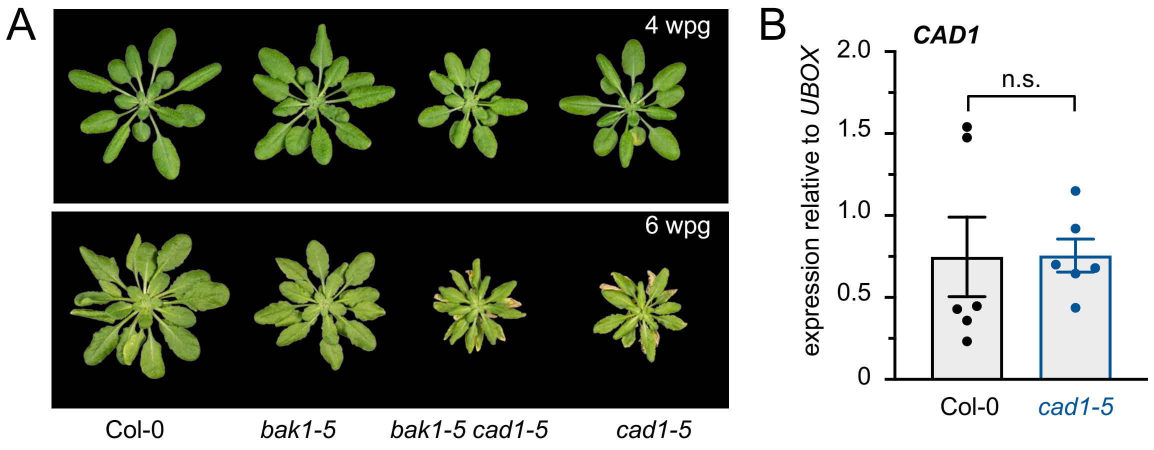


**Figure S3: *cad1-5* is not suppressed by *bak1-5*.**

**(A)** Plants were photographed after growth in a short-day chamber at 4 and 6 weeks post germination (wpg) as indicated. Genotypes were grown routinely over five years with similar phenotypes. **(B)** Real time quantitative reverse-transcription PCR of *CAD1* was performed and plotted relative to expression of *UBOX.* Values are means + standard error from six independent experiments. A paired Student's t-test indicates that there is no significant (n.s.) difference between *CAD1* expression in Col-0 and *cad1-5* (p=0.3543).

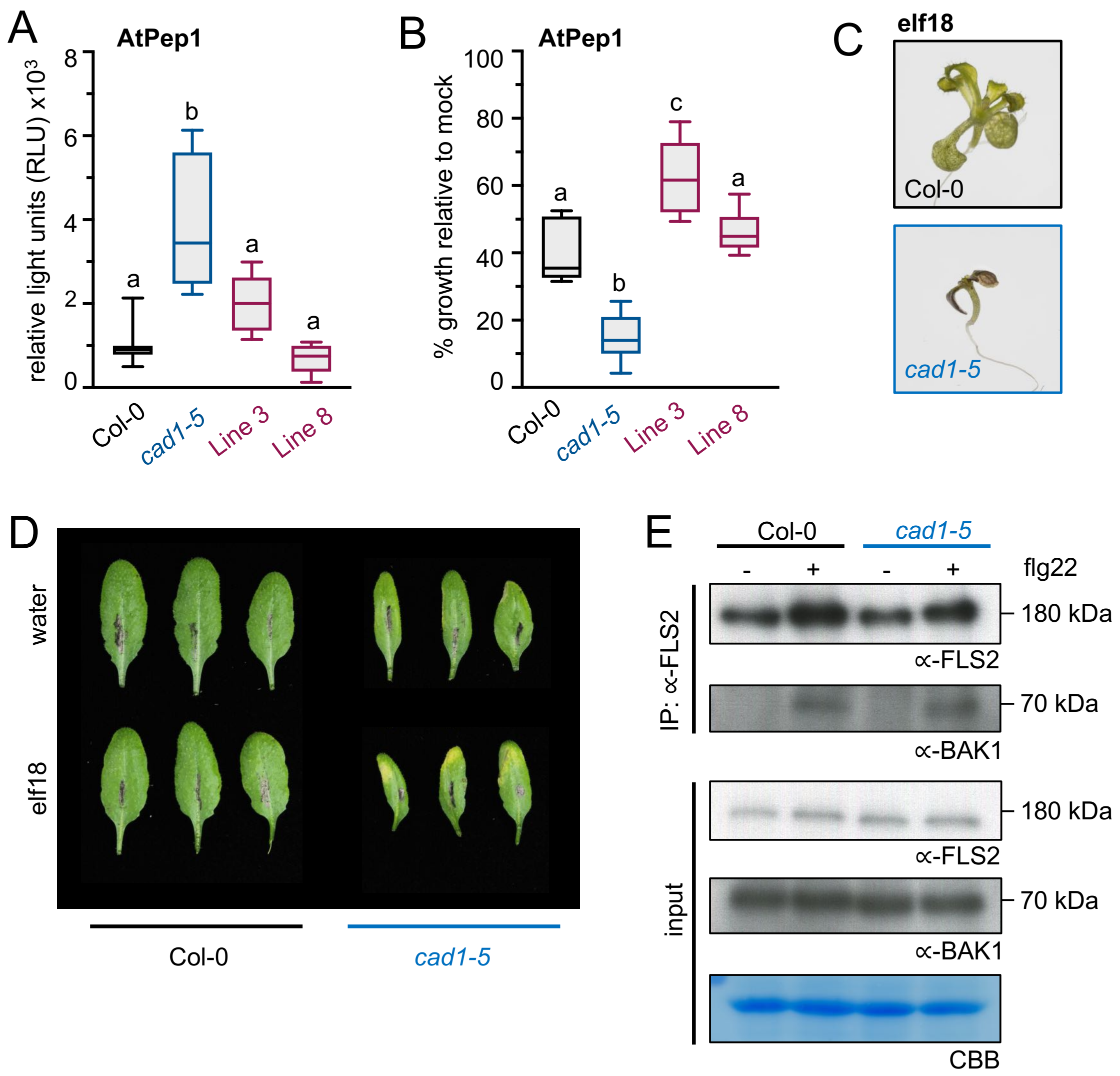


**Figure S4: Enhanced MAMP-triggered responses in *cad1-5*.**

**(A)** ROS production in Col-0, *cad1-5*, and two independent lines (Line 3 and Line 8) of *cad1-5/pCAD1:CAD1* after treatment with 1 μM AtPep1. Values represent total photon count (relative light units) over 40 min (n=6). **(B)** Seedling inhibition after 12 days of growth in 1 μM AtPep1 in the indicated genotypes, shown as % growth (fresh weight) relative to growth in control MS media (n=6). **(C)** Photograph of representative Col-0 and *cad1-5* seedlings after 12 days of growth in 100 nM elf18, indicating severe necrosis in *cad1-5*. **(D)** Five-week old plants were syringe-infiltrated with water or 1 μM elf18 and photographed after 24h. Induced cell death is observed in *cad1-5* following immune-induction. **(E)** Western blots following co-immunoprecipitation of the FLS2-BAK1 complex in Col-0 compared to *cad1-5*.
All experiments were repeated at least three times with similar results. Statistically significant ($p$<0.05) groups were analyzed by ANOVA followed by Tukey's posthoc test and are indicated by lower-case letters.

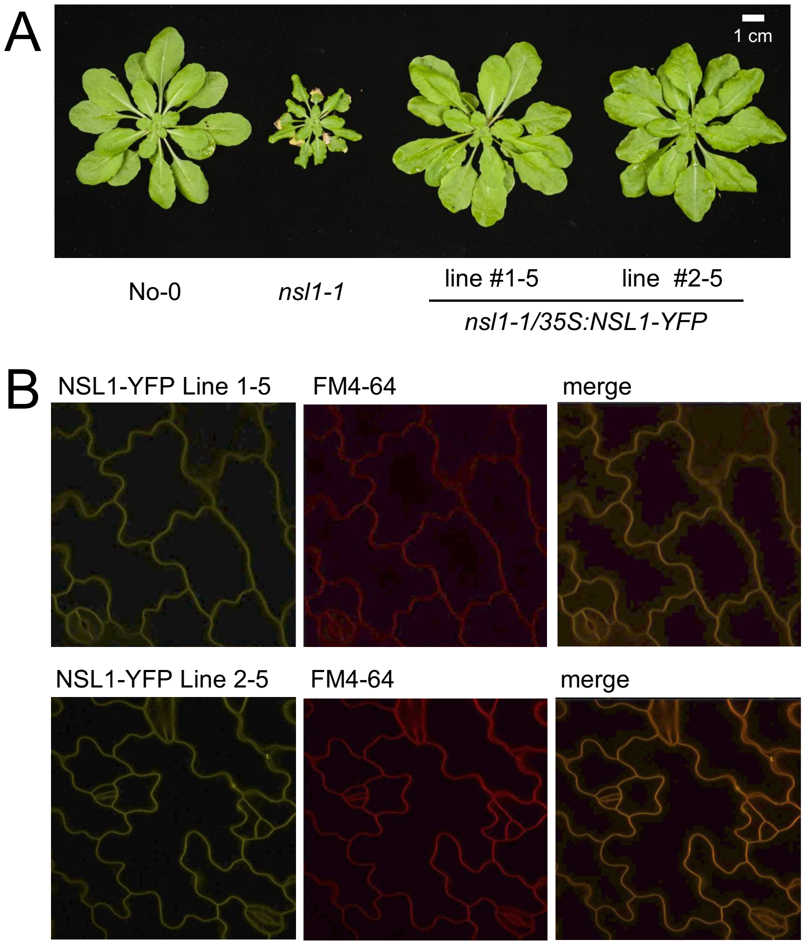


**Figure S5: NSL1 localizes to the plasma membrane.**

**(A)** Plants of the indicated genotypes (all in the No-0 ecotype) were grown in a short-day chamber and photographed 5 weeks post germination. Plants were grown several times over three years with similar phenotypes. **(B)** Confocal micrographs of NSL1-YFP expressed in cotyledon cells of two independent *nsl1-1/35S:cNSL1-YFP* transgenic lines. Seedlings were stained with the lipophilic dye FM4-64 prior to imaging to mark the plasma membrane. The merged image shows the overlay between the two channels. We observed clear plasma-membrane localization of NSL1-YFP in all four independent trials using these genotypes.

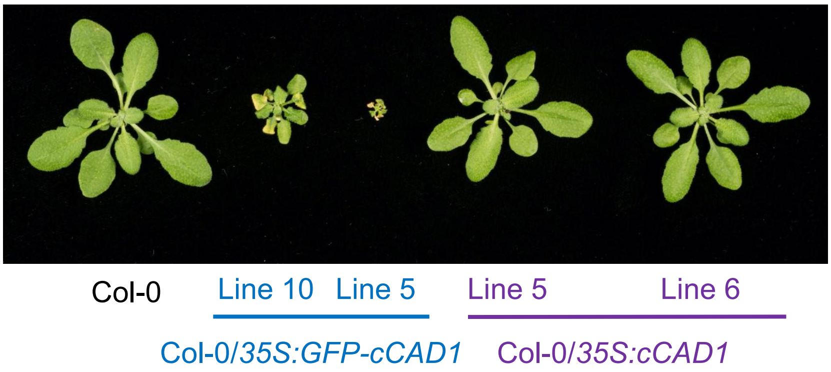


**Figure S6: N-terminally tagged CAD1 confers a dominant-negative effect.**

Plants were photographed after growth in a short-day chamber at 4 weeks post germination.

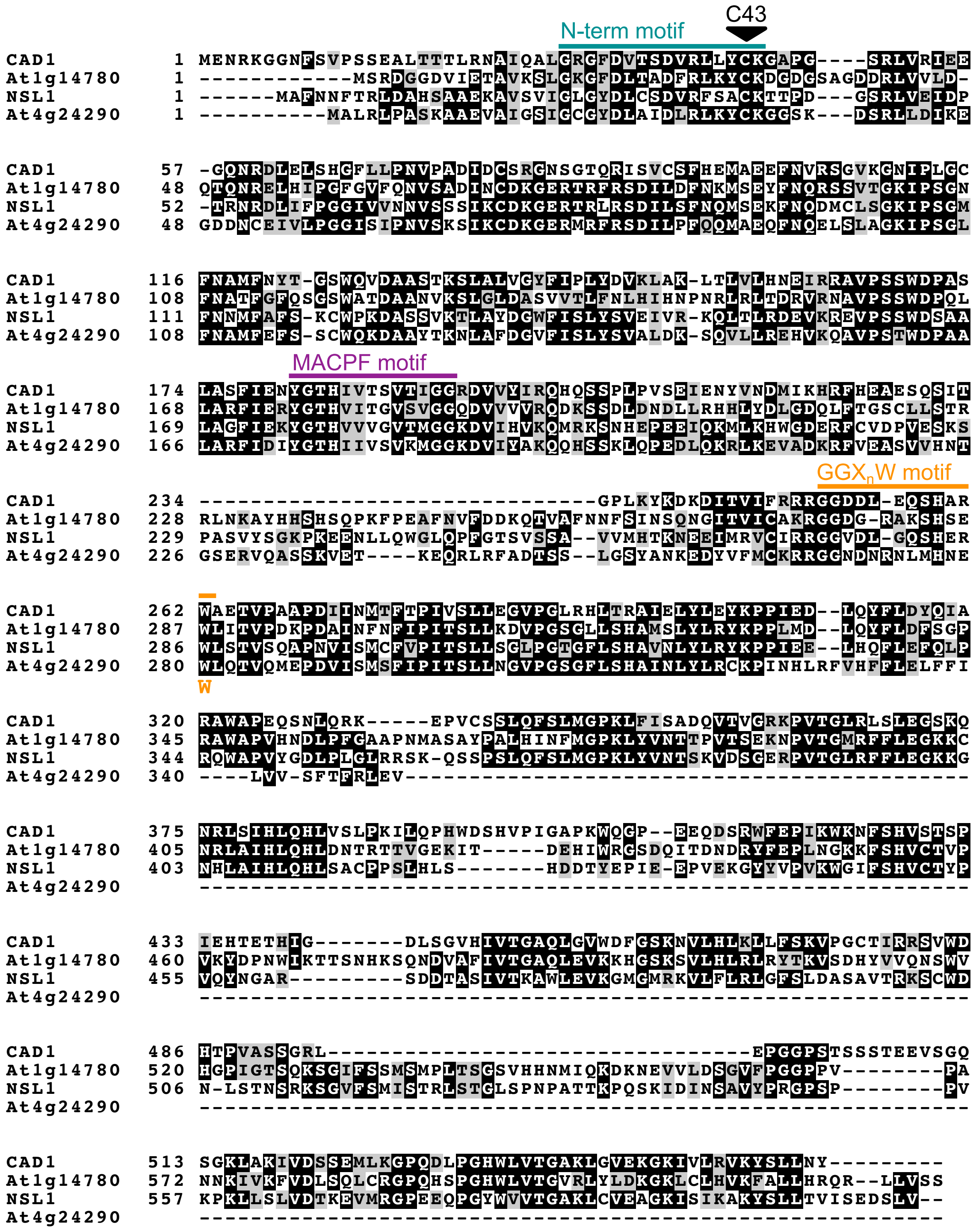


**Figure S7: Multiple sequence alignment of Arabidopsis MACPF proteins.**

Sequence alignment was performed using ClustalOmega and visualized with BoxShade. Identical residues are shown in black, similar residues in gray. The location of the conserved MACPF motif is shown in green; GGXnW motif is shown in blue; and the position of the *cad1-5* C43Y mutation is indicated by the orange arrow.

### Supplementary Table

#### Table S1: Germplasm, primers, and constructs used in this study.

| ***Arabidopsis thaliana* germplasm** | | |
|---|---|---|
| **Seed line** | **Primers used for genotyping (5'-3')** | |
| Col-0 | - | - |
| No-0 | - | - |
| *bak1-5* (Schwessinger et al., 2011) | TTGGGATCTGCAAGAGGGCTTGCGT ATTTACATGATCAGT | GAGGCGAGCAAGATCAAAAG |
| *eds1-2* (Feys et al., 2005) | ACACAAGGGTGATGCGAGACA | GTGGAAACCAAATTTGACATTAG |
| *ndr1-1* (Century et al., 1997) | GGCAACAAGCTATATGAAACC | GAATACGAGTAAATTCATCAA |
| *cad1-5* | GTGTTGGTGAGAGTCTAA | TAGACATGTCCAAAGTA |
| *cad1-2* (GK_192A09) | CTTACGCCCAACAGTTACCTG | GATCAGACGTGCTGTTCCTTC |
| *cad1-3* (GK_385H08) | TACCAAATAGCTCGAGCTTGG | TCAAACCGAACAGAACCAAAC |
| *nsl1-1* (PSH_21828) | TCCAGGTTGTTCCTCTGGAC | GTTTCAACTTCCACGCCAAT |
| *cad1-5 bak1-5* | - | - |
| *cad1-5 eds1-2* | - | - |
| *cad1-5 ndr1-1* | - | - |
| **SNP in *bak1-5 mob4*** | **Primers used for mapping (5'-3')** | |
| Chr1_1116697 | GCAGGCTCCCACCGTAA | CTGTCCAAGAGTATGACT |
| Chr1_1763961 | AGTGACATTCGGAAACC | CAAAAGTAAACATGACA |
| Chr1_5271282 | GATGCAGACCAATATT | GCAAACTGTTGAGATCT |
| Chr1_5276811 | TTGACCCTCCTTCCCAGCCTC | CTCTTAGATTGTGCCTTGT |
| Chr1_5740226 | ATGAGCCAGCAGGTATGA | CACCAGTTACTGCAGCTTTGT |
| Chr1_5741941 | TATCTTCTTATACCAAG | CTGCTAGTTTGACCACAGT |
| Chr1_5901236 | CTCGAGCTTTCTCATAA | ACTATGCTGCCGTGATCTTGCTG |
| Chr1_6646943 | GTCTGGAAGTTTCTAGAG | TGCGTTAAAGCCAGTCAACCG |
| Chr1_6681734 | GAGTCAGATGATAGAGAC | GAGTAACTTCACCATATTC |
| Chr1_6934331 | GTATCGAGAGTGAATTCACT | GCTCCATTAGATGGAGACTTC |
| Chr1_7073500 | ACGACCCGAGTAGTAATGCT | GCAGTCAGCGACGAAGCCAAAGAT |
| Chr1_8351039 | CTCGCACTTTCTTCTAGACTC | GAGATGATTCTTTGAATCT |
| Chr1_9042744 | GGGGAGAAGATTCTGGTG | GTTAGAATCTGGTCTTGG |
| Chr1_9194612 | AAGATGTTCAATGTGAAGAAT | GCTTTCTTAACCATTG |

| | | |
|---|---|---|
| Chr1_9437278 | CAGGTGGATGAGAATTTG | CCACCGCGCTTCCGTTGAT |
| Chr1_9525661 | TCTGTGGTTAGCAATGT | ACAGACTGTTTCCACC |
| Chr1_10381734 (*cad1-5*) | GTGTTGGTGAGAGTCTAA | TAGACATGTCCAAAGTA |
| Chr1_10752574 | ATTCCTCTTGCAGTTTAATAG | TGATATCAACGCTTGAGGTTTC |
| Chr1_10815851 | AGATGTGAGTATGGAAG | GACTCCATTATCCATCATC |
| Chr1_11969678 | CATATCGCTTGTAGATGAACT | TTCTCATAAGAGTCTGGCTTCTGG |
| Chr1_12375231 | TTGGAGCTGGAGCCACA | GTCACCCGAATCTTGA |
| Chr1_13774328 | GAATGTCTTGAGAGGCGATCT | CTATGCCTTAGATGAATGCAGA |
| Chr1_14051638 | GCATTGATCCGGCTACATTG | GCAGTCATTCAAGGTCGATG |
| Chr1_18510852 | GCATTACTTGTGAAGTCAAGT | TCTTCATATTCGACAACATCCTTC |
| Chr1_18872523 | TCGGATGAGCAGCTGAAGCAGC | TAGATTCGTCAACTCAGGA |
| Chr3_1308089/1217 | TCTCAAGATTACCGTAATCAG | ATCCAACTCTGCATCTCCTTGT |
| Chr4_7211101 | ATGATAATTGTCCAATCGA | GTTGATCCAAACCTTGGCTTGAAC |
| Chr5_5599873 | GAAGACAGTTGTGAACAAGTTG | GCACTGTACGTATCACAACTTG |
| Chr5_9186001 | GTGAGCCTGAAGGAGATGTT | TGGAATCCCTGTCTGACACAATA |
| Chr5_20881895 | AATCTTGTTTCATTTGAGC | TAATACCTACCTCAACGCT |
| Chr5_21236328 | CACACCACAAATTTCGA | CAGTCTGAGCTCCCAAGACGG |

| **Seed line** | **Construct used** | ***In planta* resistance marker** |
|---|---|---|
| *bak1-5 mob4/ pCAD1:gCAD1 line 7* | pGWB1 - *pCAD1:gCAD1* | Kanamycin |
| *bak1-5 mob4/ pCAD1:gCAD1 line 9* | | |
| *cad1-5/ pCAD1:gCAD1 line 3* | | |
| *cad1-5/ pCAD1:gCAD1 line 8* | | |
| *cad1-5/ pCAD1:gCAD1-GFP line 4* | pGWB4 - *pCAD1:gCAD1-GFP* | Kanamycin |
| *cad1-5/ pCAD1:gCAD1-GFP line 5* | | |
| *Col-0/ 35S:GFP-gCAD1 line 10* | pGWB6 - *35S:GFP-gCAD1* | Kanamycin |
| *Col-0/ 35S:GFP-gCAD1 line 5* | | |
| *Col-0/ 35S:gCAD1 line 5* | pGWB2 - *35S:gCAD1* | Kanamycin |
| *Col-0/ 35S:gCAD1 line 6* | | |
| *nsl1-1/ 35S:NSL1-YFP line 1* | pXCSG - *35S:NSL1-YFP* | Basta |
| *nsl1-1/ 35S:NSL1-YFP line 2* | | |

| Gene constructs | | |
|---|---|---|
| **Entry vectors** | **Primers used for cloning (5'-3')** | |
| pENTR - *cCAD1*<br>(no promoter, no stop codon) | CACCATGGAGAATCGTAAAGGAGG | ATAATTTAGCAACGAATACTT |
| pENTR - *cCAD1*<br>(no promoter, native stop codon) | CACCATGGAGAATCGTAAAGGAGG | TCAATAATTTAGCAACGAATA |
| pENTR - *gCAD1*<br>(native promoter, native stop codon) | CACCCATCCTCTAATTTGAAGCAT | TCAATAATTTAGCAACGAATA |
| pENTR - *gCAD1*<br>(native promoter, no stop codon) | CACCCATCCTCTAATTTGAAGCAT | ATAATTTAGCAACGAATACTT |
| pENTR - *gCAD1*<br>(no promoter, no stop codon) | CACCATGGAGAATCGTAAAGGAGG | ATAATTTAGCAACGAATACTT |
| pENTR - *gCAD1-C43Y*<br>(no promoter, no stop codon)<br>*amplified from *cad1-5* gDNA | CACCATGGAGAATCGTAAAGGAGG | ATAATTTAGCAACGAATACTT |
| pENTR - *cNSL1*<br>(no promoter, no stop codon) | CACCATGGCCTTTAACAATTTCAC | GACCAGTGAGTCTTCCGATA |
| **Destination vectors** | **Entry vector** | **Backbone** |
| pGWB1 - *pCAD1:gCAD1* | pENTR - *gCAD1*<br>(native promoter, native stop codon) | pGWB1 |
| pGWB4 - *pCAD1:gCAD1-GFP* | pENTR - *gCAD1*<br>(native promoter, no stop codon) | pGWB4 |
| pGWB6 - *35S:GFP-gCAD1* | pENTR - *cCAD1*<br>(no promoter, native stop codon) | pGWB6 |
| pGWB2 - *35S:gCAD1* | pENTR - *cCAD1*<br>(no promoter, native stop codon) | pGWB2 |
| pXCSG - *35S:gCAD1-YFP* | pENTR - *gCAD1*<br>(no promoter, no stop codon) | pXCSG |
| pXCSG - *35S:gCAD1-C43Y-YFP* | pENTR - *gCAD1-C43Y*<br>(no promoter, no stop codon) | pXCSG |
| pXCSG - *35S:NSL1-YFP* | pENTR - *cNSL1*<br>(no promoter, no stop codon) | pXCSG |

| Primers used for quantitative real-time PCR | | |
|---|---|---|
| **Target gene** | **Primers used for qPCR (5'-3')** | |
| *PR1* | GTAGGTGCTCTTGTTCTTCCC | CACATAATTCCCACGAGGATC |
| *UBOX* | TGCGCTGCCAGATAATACACTATT | TGCTGCCCAACATCAGGTT |
| *CAD1* | GCTTCCATGAGATGGCAGAA | CAATAAAGCTAGCTAGGGAAGC |
